# Fast analysis of Spatial Transcriptomics (FaST): an ultra lightweight and fast pipeline for the analysis of high resolution spatial transcriptomics

**DOI:** 10.1101/2024.07.30.605511

**Authors:** Valerio Fulci

## Abstract

Recently, several protocols repurposing the Illumina flow cells as an RNA capture device for spatial transcriptomics have been reported. These protocols yield high volumes of sequencing data which are usually analyzed through the use of HPC clusters. I report inhere a novel pipeline for the analysis of high resolution spatial transcriptomics datasets obtained on Illumina flow cells. FaST is compatible with OpenST, seq-scope and potentially other protocols. It allows full reconstruction of the spatially resolved transcriptome, including cell segmentation, of datasets consisting of more than 500 M million reads in as little as two hours on a standard multi core workstation with 32 Gb of RAM. The FaST pipeline returns RNA segmented ST datasets suitable for subsequent analysis through commonly used packages (e.g scanpy or seurat).

Notably, the pipeline I present relies on the spateo-release package for RNA segmentation, and does not require Hematoxylin/Eosin or any other imaging procedure to guide cell segmentation. Nevertheless, integration with other software for imaging-guided cell segmentation is still possible.

## Introduction

Spatially resolved transcriptomics (ST) has recently emerged as a powerful technique to investigate the spatial patterns of gene expression in tissues, in physiologic as well as pathologic contexts [1]. While the earliest approaches yielded gene expression information with a medium resolution (about 100 μm), more recent protocols proved able to faithfully record spatially resolved gene expression at submicrometric resolution, allowing analysis of these datasets at cellular and subcellular level. In particular, a series of recent works has independently reported that Illumina flow cells may be used as an RNA capture device to yield ST datasets at a resolution of about 0.6 μm, closely resembling the resolution power of optical microscopy and faithfully recapitulating the expected histological features of the analyzed tissues [2–4]. A typical ST experiment on a 10 to 15 mm^2^ specimen will yield 0.5 to 1 billion reads posing a considerable challenge to rapid and efficient downstream analysis. Importantly, Illumina flow cells are organized as an array of “tiles” each with a surface of about 1 mm^2^. A typical ST experiment will involve 15 to 20 such “tiles”. While recently published papers mainly focus on “proof of principle” specimens, with a deep focus on a single or few samples, unleashing ST potential will require analysis of large cohorts of samples, suggesting the need for a lightweitìght, reproducible and fast analysis pipeline that could be routinely applied in both small scale and large scale dataset to provide a univocal method for Illumina flow cells based ST datasets.

One of the major issues posed by analysis of high resolutions ST is represented by the cell segmentation step of the analysis. Earliest approaches have leveraged tissue staining techniques highlighting cell nuclei (e.g. DAPI, H&E) thus providing compelling evidence that the density and identity of the captured RNAs faithfully mirrors the composition of the analyzed tissues at single cell level [2,4]. However, such approaches require tissue staining, image acquisition and image data analysis, resulting in complicated and poorly scalable protocols both in terms of wet lab and bioinformatic analysis. Indeed, several recent bioinformatic approaches have been developed to perform segmentation of ST data analysis into putative single cells guided in part or completely by RNA density as assessed by ST [5–8].

Inhere, I describe the Fast analysis of ST (FaST) pipeline, a simple yet rigorous algorithm with a low memory footprint, which allow very quick analysis of large ST datasets yielding RNA segmented datasets in anndata [9] formats suitable for downstream analysis with widespreadly used single cell transcriptome data analysis tools [10,11].

The data presented inhere show that RNA segmented “pseudo cells” recapitulate faithfully the results obtained by image based cell segmentation, both in terms of cell type identification, cell number and specificity of gene expression.

FaST has been written largely in bash and perl and by design has a minimal set of requirements, to ensure long term reliability with minimal maintenance of the pipeline.

## Methods

### Flowcell barcode map preparation

FaST reads the HDMI fastq file obtained in the first round of sequencing and outputs a “flow cell barcode map” consisting of a set of files (one for each tile in the flow cell) each listing the HDMI barcodes associated with x and y coordinates as reported in the header of the reads. If identical barcodes occur within a tile, those are discarded. Identical barcodes across different tiles are kept and later handled during sample analysis.

At the same time, an index consisting of a random sample of the barcodes from each tile is saved to allow fast retrieval of tiles in each experiment during data analysis.

### Sample fastq reads preprocessing

FaST reads the whole set of R1 reads (containing spatial barcodes). To identify the tiles of the Illumina flow cell used for RNA capture, FaST compares the whole set of R1 reads with the Illumina flowcell barcode map index generated with FaST-map, retaining tiles for which at least 10% of the indexed barcodes are present in the R1 file.

Following the identification of the tiles used in the experiment, only barcodes mapping to these tiles are retained, and all reads corresponding to a different barcode are discarded. Barcodes mapping to multiple tiles in the flow cell, but mapping to one and only one barcode within the selected tiles are retained. Any other ambiguous barcode (i.e., matching more than one barcode in the selected tiles) will be discarded.

Following the R1 reads analysis and barcode selection, R2 reads will be collected and converted into a unaligned BAM file with the following attributes as BAM tags:

R1 barcode: BC:Z: tag

Tile name: RG:Z:tag

UMI: MI:Z:tag

X coordinate offset within tile: CX:i:tag

Y coordinate offset within tile: CY:i:tag

A (temporarily) empty (NA) putative subcellular localization tag (LC:Z:NA).

### Reads alignment

FaST uses a single alignment step with STAR [12]. The STAR index is generated by the FaST-reference script, which first hard-masks (i.e. converts into “N”) nucleotides in all regions of the genome outside of the chosen reference gtf. This choice is motivated by the fact that DGE algorithms retain any aligned read which has one and only one valid alignment within the transcriptome, regardless of the number of “intergenic” alignments observed. It is therefore pointless to compute whether there are or not alignments in the non-coding regions of the genome. This has a twofold advantage:

1. The size of the effective STAR genome index is consequently significantly smaller.
2. Masking of highly repeated rRNAs loci (which are not part of the Gencode annotations) allows to concatenate the 45S DNA region and PhiX genomes as two extra chromosomes to the genome fasta, allowing to map rRNAs and PhiX in a single step together with transcriptome.

During alignment polyA tail will be clipped by STAR. All BAM tags will be retained and reads with a valid alignment will be output in BAM format.

### Digital Gene Expression

The BAM file is then split for parallel processing tile by tile, piping reads belonging to the same tile (based on the RG:Z: tag of the read) to parallel processes. Each process will:

1. Remove and count reads mapping to rRNA and PhiX (selected based on the chromosome names) to output PhiX and rRNA mapping reads statistics.
2. Convert the SAM format alignment to a BED format, retaining BAM tags as a 7th field of the BED entry. Alignments will be split on CIGAR N operations, in this latter case the subcellular localization tag will be set to LC:Z:CY (for cytoplasmic).
3. Using bedtools intersect [13], overlaps of each BED entry with a custom BED annotation (generated by the FaST-reference script) are computed.
4. A custom perl script parses genomic intervals overlapped by each read, setting the subcellular localization to cytoplasmic (LC:Z:CY) if the gene is a mitochondrial gene and to nuclear if the read overlaps any intron of the gene body (LC:Z:NU). Due to the lack of introns in mitochondrial genes, these two classifications will be necessarily mutually exclusive for reads mapping to a single gene body. If all mappings of the same read map to the same gene body, an output will be generated, consisting of the gene name, the barcode, the UMI and the x and y coordinates of the barcode and the putative subcellular localization.
5. The output of step 3 will be piped to GNU sort and to GNU uniq commands, yielding gene mappings with unique UMIs and barcodes.
6. A custom perl script loads all the data of each tile into a sparse “barcodes x gene counts” matrix and writes MatriMarket format output and a text file in the spateo-release input format.

### RNA based cell segmentation

Cell segmentation starts with a list of nuclear localized transcripts including microRNA host genes and APEX-seq [14] determined nuclear transcripts to obtain a putative nuclear mask applying the EM+BP algorithm. Then the intron counts are used to perform a second round of segmentation and resulting masks are joined with those obtained in the previous step. Finally, whole cells are segmented using the entire matrix of all reads and joined with the masks obtained in the previous step. Segmented cell counts and spatial coordinates are exported in Anndata format. Finally, the anndata [9] objects (one for each tile) will be concatenated into a single anndata file, in this step the coordinates of each tile are corrected according to the appropriate offsets to rebuild the full image of the sample.

Alternatively, users can opt to use a data driven approach entirely based on the sequencing data. Briefly, pucks which contain at least 10 UMIs will be assigned a “cytoplasmic” and a “nuclear” score, based on the number of spliced and mitochondrial reads (cytoplasmic score) or miRNA host genes and intronic reads (nuclear score). For each tile, counts assigned to the pucks attaining a cytoplasmic score above the 90th percentile of the overall cytoplasmic score will be collected as “cytoplasmic” counts for that tile. An analogous approach will lead to the collection of the “nuclear” counts for each tile. These count files are fed to differential expression analysis with DESeq2 [15] to yield a list of predominantly nuclear genes to be used in the first segmentation step.

### Benchmarking

Benchmarks were performed either in a “workstation” or in an “HPC” hardware. The “workstation” hardware is equipped with a 12th Gen Intel® Core™ i9-12900K processor (16 CPU cores, 8 of which are multithreading cores, for a total of 24 threads) and 32Gb of RAM. All jobs were launched with a 24 threads setting in this hardware.

A single node of the HPC hardware used for benchmarking is equipped with 2 AMD EPYC™ 7452 processors (32 CPUs, 64 threads each) and 256 Gb of RAM. All jobs were launched with a 32 threads setting in this hardware. This “HPC” enforces a rule making available no more than 2 Gb of RAM for each thread, resulting in a maximum of 64Gb of RAM when 32 threads are reserved through slurm. To ensure that spacemake and NovaScope jobs requiring more than 64 Gb of RAM were not killed, a considerably higher number of threads was reserved, although the actual jobs were launched on no more than 32 threads. Due to the limited memory footprint of FaST, this was not needed when running FaST. The option -H of FaST triggers parallel pre-processing of pairs of fastq files (when the dataset consists of more than one pair fastq files, resulting in a considerably higher memory footprint in the HPC setting compared to the workstation.

### Downstream analysis

Anndata object were analyzed with scanpy 1.10.1 [11], filtering with identical parameters (min counts/cell=200; max counts/cell = 2000, mitochondrial counts % < 20, minimum number of cells expressing a gene to retain it= 10) for all datasets. Data were normalized and log transformed. Leiden algorithm was used for clustering.

### Data and code

OpenST data are available through GEO (GSE251926) NovaScope data were obtained from this website https://seqscope.github.io/NovaScope/getting_started/access_data/). To obtain the 1.8 Billion reads dataset the following samples were pooled together: N3_B08C_v2b; N3_B08C_v2bu; N3_B08C_v2a; N3_B08C_v2cu.

FaST is available on github: https://github.com/flcvlr/FaST

Genomic reference files were downloaded from GENCODE[16]. Version 46 (GRCh38) was used for human samples; version M35 (GRCm39) was used for mouse samples. Human (URS0000ABD7E8_9606) and mouse (URS00026C81C6_10090) 45S rRNA sequences were obtained from RNA central[17]. PhiX genome sequence (NC_001422.1) was obtained from NCBI Nucleotide.

## Results

### The FaST pipeline

FaST is mainly written in perl and bash and has a very limited set of dependencies (STAR, samtools, bedtools and the python packages spateo-release and scanpy). The first steps of the pipeline consist in the collection of the spatial barcodes of the experiment (“read 1”), the selection of the “tiles” of the Illumina flow cell corresponding to that set of barcodes and the mapping of the reads to the reference genome with STAR. Additionally, several data, including spatial coordinates, putative subcellular localization, UMI and barcode are attached to each read as BAM tags, thus avoiding time consuming downstream steps to re-match barcode, spatial coordinates and read alignment data (Figure 1).

**Figure 1:**
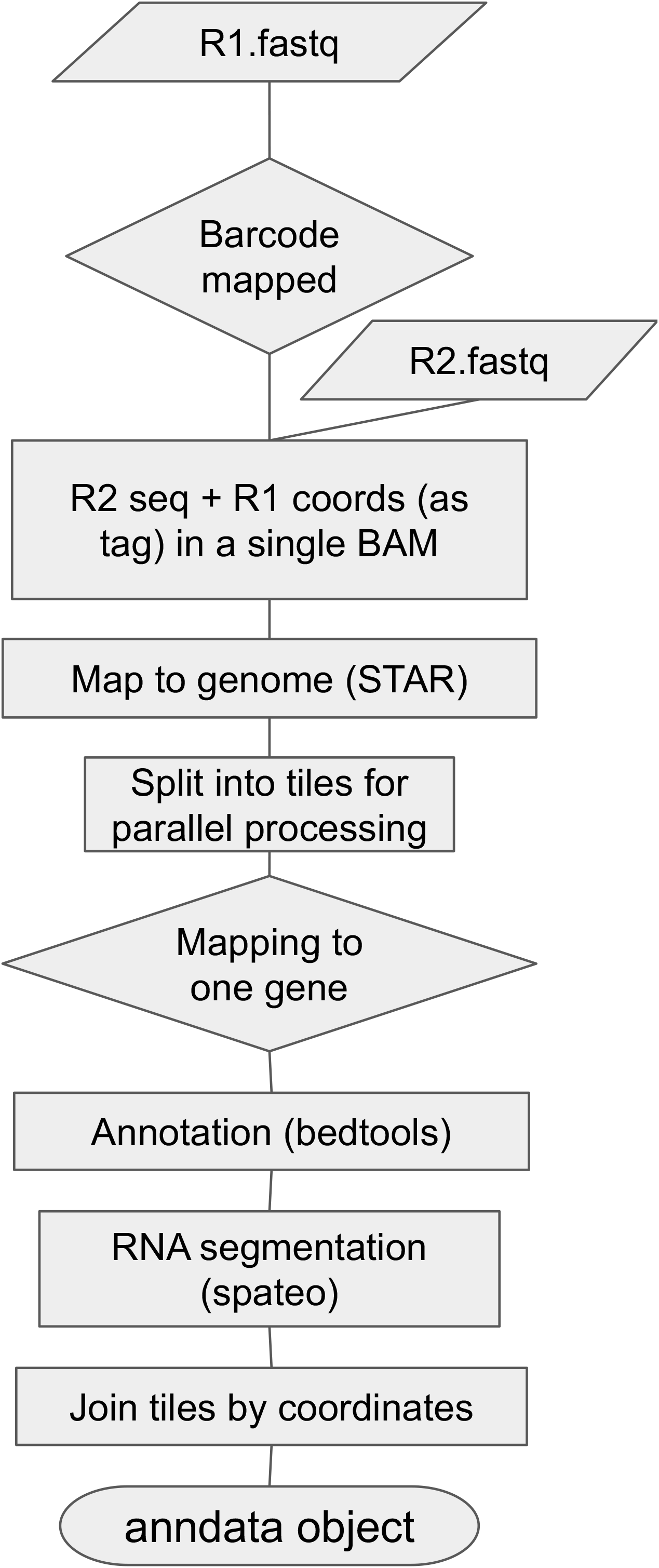
Flowchart representing the major steps of the FaST pipeline.

Downstream of reads alignment on the reference genome using STAR [18], reads are parsed by custom scripts which are launched in parallel on each tile of the Illumina flow cell used in the experiment. Reads are first compared with Gencode annotation [16] and checked for overlap with one and only one gene. Reads spanning an exon/exon junction or a mitochondrial gene are labeled as cytoplasmic, while reads overlapping intron sequences or mapping to microRNA host genes are labeled as nuclear. A further tag corresponding to the gene is added in this step.

Reads mapping to the same gene and carrying the same barcode and UMI are collapsed and used to build a “puck”-level sparse count matrix for each tile. This data can be easily imported into Seurat, Scanpy or other similar tools and converted in further formats to be fed to a variety of downstream cell segmentation packages, including OpenST.

The sparse matrix is then used by FaST to generate the input file for the spateo package for each tile. RNA segmentation is performed on each tile separately and the RNA segmented tiles are finally rejoined into a single anndata object.

RNA segmentation leverages a pool of nuclearly localized transcripts, as observed through Apex-seq [14]. However, an option to run discovery of nuclear transcripts based on the colocalization at puck level with intronic reads and microRNA precursors is also available.

### FaST can quickly process large amounts of data with minimal hardware requirements

Aiming to obtain a lightweight software for the analysis of high resolution ST repurposing Illumina flowcells as RNA capture devices, I benchmarked my pipeline on a small workstation equipped with a 12th Gen Intel® Core™ i9-12900K processor (16 CPU cores, 8 of which are multithreading cores, for a total of 24 threads) and 32Gb of RAM. Currently, such workstations can be purchased for around 1000 Euros each. To validate my pipeline I chose to analyze the samples listed in Table 1.

**Table 1:** List of datasets used to validate FaST with overview of time requirements on a workstation*.

| Dataset | Protocol | GEO # | Reads (M) | Wall time (mapping + DGE) | Wall time (segmentation ) | Wall time (full analysis) |
| --- | --- | --- | --- | --- | --- | --- |
| Mouse head e13 | OpenST | GSM7990097 | 569.6 | 0:43:39 | 0:55:40 | 1:39:19 |
| Metastatic Lymphnode S4 | OpenST | GSM7990102 (SRR27331444) | 782.7 | 1:12:31 | 1:18:57 | 2:31:28 |
| Mouse liver | Seq-scope | N/A | 1781.9 | 3:54:07 | 1:32:01 | 5:26:08 |

\* All these tests were carried out on a workstation equipped with an 12th Gen Intel® Core™ i9-12900K and 32Gb of RAM

FaST could successfully analyze both sample GSM7990097 (596 millions reads; elapsed wall time: 1 hour 40 minutes) and GSM7990102 (781 Million reads; elapsed wall time 2 hours 31 minutes).

To ascertain if the limited hardware could also accommodate analysis of significantly larger datasets, I also tested FaST on the recently published NovaScope dataset [3], finding that FaST can handle equally well these data (∼1.8 Billion reads, datasets N3_B08C_v2b, N3_B08C_v2bu N3_B08C_v2c, N3_B08C_v2cu) with an elapsed wall time 5 hours and 26 minutes and a memory footprint of 27.8 Gb (Table 2).

**Table 2:** Benchmarking data for preliminary Tasks performed by FaST on a workstation*.

| Task | Fastq file size (Gb) | Wall time (mm:ss) | User time (seconds) | %CPU | RAM (max, Gb) |
| --- | --- | --- | --- | --- | --- |
| Extract barcode coordinates for entire flowcell | 400 | 3:21:50 | 24125.16 | 224% | 0.005760 |
| Generate reference (hg38) | N/A | 14:08 | 9817.73 | 1170% | 26.9 |
| Generate reference (mm39) | N/A | 9.44 | 5263.40 | 911% | 26.9 |

\* All these tests were carried out on a workstation equipped with an 12th Gen Intel® Core™ i9-12900K and 32Gb of RAM
\*\* This task is performed only once to obtain a barcode map of an entire S4 flowcell, sufficient to perform at least 100 different experiments.

To compare FaST with existing alternative pipelines, I compared the performance of FaST, NovaScope[3] and spacemake[19] in an HPC environment. This comparison has been limited to the mapping step, as neither spacemake nor NovaScope perform cell segmentation of data, which is performed instead by the OpenST[4] python package or the Ficture[6] package, respectively.

In an HPC environment FaST proved to run slightly less fast (this is expected, as single processors on HPC tend to have a lower frequency compared to small workstations), with a wall time of 1 hour and 44 minutes for sample GSM7990097 (memory footprint of 23.4 Gb), 1 hour and 44 minutes for seq-scope dataset N3_B08C_v2cu (memory footprint of 23.2 Gb) and 2 hours and 25 minutes for the mapping of sample GSM7990102, (memory footprint of 28.9 Gb).

On the one hand, NovaScope required, for sample N3_B08C_v2cu, 2 hours and 46 minutes (memory footprint 28.7 Gb). Larger datasets (Table 3) up to 1.3 Billion reads highlight a roughly linear increase in the processing time for NovaScope (Figure S1 and Figure S2), while FaST, in an HPC environment, scales quite well, provided that data are split in independent fastq files.

**Table 3:**
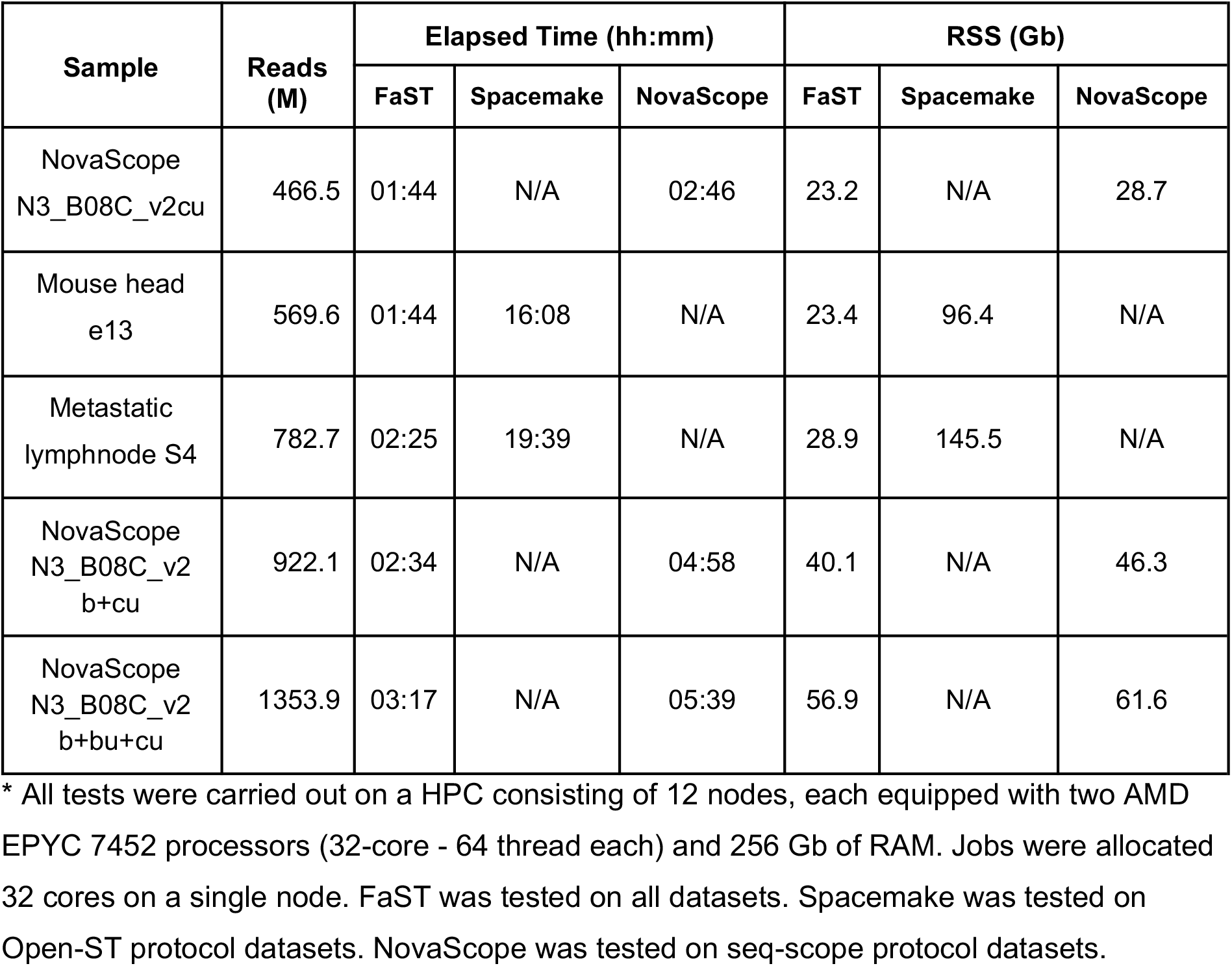
Comparison with existing alternatives on HPC (mapping + SGE)

| Sample | Reads (M) | Elapsed Time (hh:mm) |  |  | RSS (Gb) |  |  |
| --- | --- | --- | --- | --- | --- | --- | --- |
|  |  | FaST | Spacemake | NovaScope | FaST | Spacemake | NovaScope |
| NovaScope N3_B08C_v2cu | 466.5 | 01:44 | N/A | 02:46 | 23.2 | N/A | 28.7 |
| Mouse head e13 | 569.6 | 01:44 | 16:08 | N/A | 23.4 | 96.4 | N/A |
| Metastatic lymphnode S4 | 782.7 | 02:25 | 19:39 | N/A | 28.9 | 145.5 | N/A |
| NovaScope N3_B08C_v2 b+cu | 922.1 | 02:34 | N/A | 04:58 | 40.1 | N/A | 46.3 |
| NovaScope N3_B08C_v2 b+bu+cu | 1353.9 | 03:17 | N/A | 05:39 | 56.9 | N/A | 61.6 |

\* All tests were carried out on a HPC consisting of 12 nodes, each equipped with two AMD EPYC 7452 processors (32-core - 64 thread each) and 256 Gb of RAM. Jobs were allocated 32 cores on a single node. FaST was tested on all datasets. Spacemake was tested on Open-ST protocol datasets. NovaScope was tested on seq-scope protocol datasets.

On the other hand, spacemake required 96 Gb of RAM to successfully carry out mapping of sample GSM7990097 with a wall time of 16 hours and 145.5 Gb for sample GSM7990102, with a wall time of 19 hours and 40 minutes (Figure S1 and Figure S2 S2). I did not test spacemake on larger datasets.

### FaST analysis yields the expected number of cells and the expected patterns within tissue

Full analysis of dataset GSM7990097 (open ST protocol, mouse e13 head) consisting of 596 million raw reads) with the FaST pipeline yielded a total of 54437 cells, with a mean of 571 UMIs per cells, accounting for 31.1 Millions UMIs (Fig 2). The overall organization is extremely similar to the one reported in the GEO deposited data through H&E segmentation, which yielded a comparable number of cells (49043). The UMAP projection of the dataset, although the arrangement of the clusters in the UMAP space is different, suggests that the overall tissue organization is correctly represented (Fig 2). In particular, in both plots the three major clusters identified in the forebrain (clusters 0 and 3 in panels A and C; clusters 1, 2 and 7 in panels B and D), have a similar distribution in the UMAP space.

**Figure 2:**
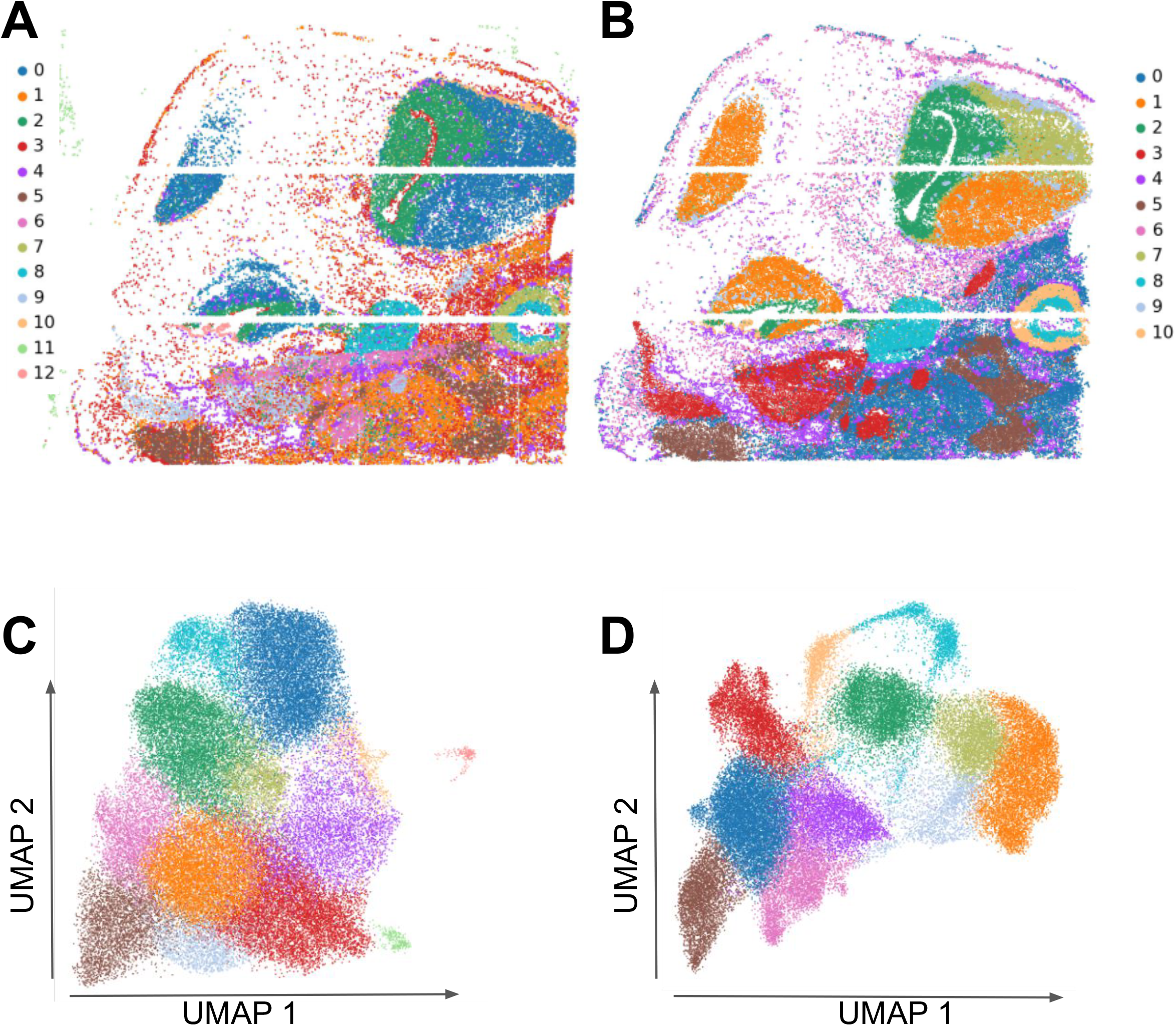
Analysis of sample GSM7990097 (mouse embryo head) using FaST (panels A and C) compared to GEO deposited processed data. Panel A: Spatial gene expression was reconstructed using FaST. Segmented gene expression in anndata format was used to cluster “pseudo-cells” using scanpy with default parameters. Clustered cells were plotted in spatial coordinates. Panel B: Segmented gene expression in anndata format (GEO) was used to cluster cells using scanpy with default parameters. Clustered cells were plotted in spatial coordinate. Panel C: UMAP plot of pseudo-cells obtained using FaST. Panel D: UMAP plot of segmented cells from GEO processed data (GSM7990097). Cluster colors do not correspond to the same cell types in the two different protocols. Colors were automatically assigned based on the abundance of each cluster in each output (FaST, panels A and C; Spacemake/OpenST panels B and D).

As an example of a human sample, I chose the section 4 of a metastatic lymph node (GEO GSM7990102). My pipeline yielded a total of 91304 cells, which included 61.8 M UMIs, with an average of 677 UMIs per cell (Fig 3). We compared our findings with the H&E segmented dataset deposited on GEO in anndata format. Applying the identical thresholds for analysis, H&E segmentation yielded 68423 cells on the same sample, with a mean of 848 counts/cell, accounting for 58 M UMIs. In this case FaST identified a significantly higher number of cells, most of which are localized in the cancerous part of the lymphnode (metastatic tumor cells, clusters “0” and “7” in figure 3 Panel A). In this case a slightly better resolution seems to be achieved by FaST, which clearly identifies keratin pearls (cluster 7, Figure 3A) as a specific cell type. On the other hand, the Spacemake/OpenST protocol only identifies a single major population of cancerous cells (cluster 1, Fig 3B). However, the spacemake/OpenST protocol seems to capture a higher diversity in the lymphoid part of the tissue (clusters 1, 4 and 8 in Fig 3 A; clusters 0, 4, 5, 6 and 7, Fig 3B). UMAP projections (Fig 3 panels C and D) confirm that both protocols yield a neat segregation of epithelial cancer cells (panel C:upper left, clusters 2 and 7; panel D: right side, cluster 2) vs lymphoid cells. Plots of individual genes (Figure S3) highlight the overall agreement in the spatial pattern of gene expression between FaST and OpenST and confirm the above mentioned differences in the segmentation of the cancerous tissue and the lymph node between the two algorithms..

**Figure 3:**
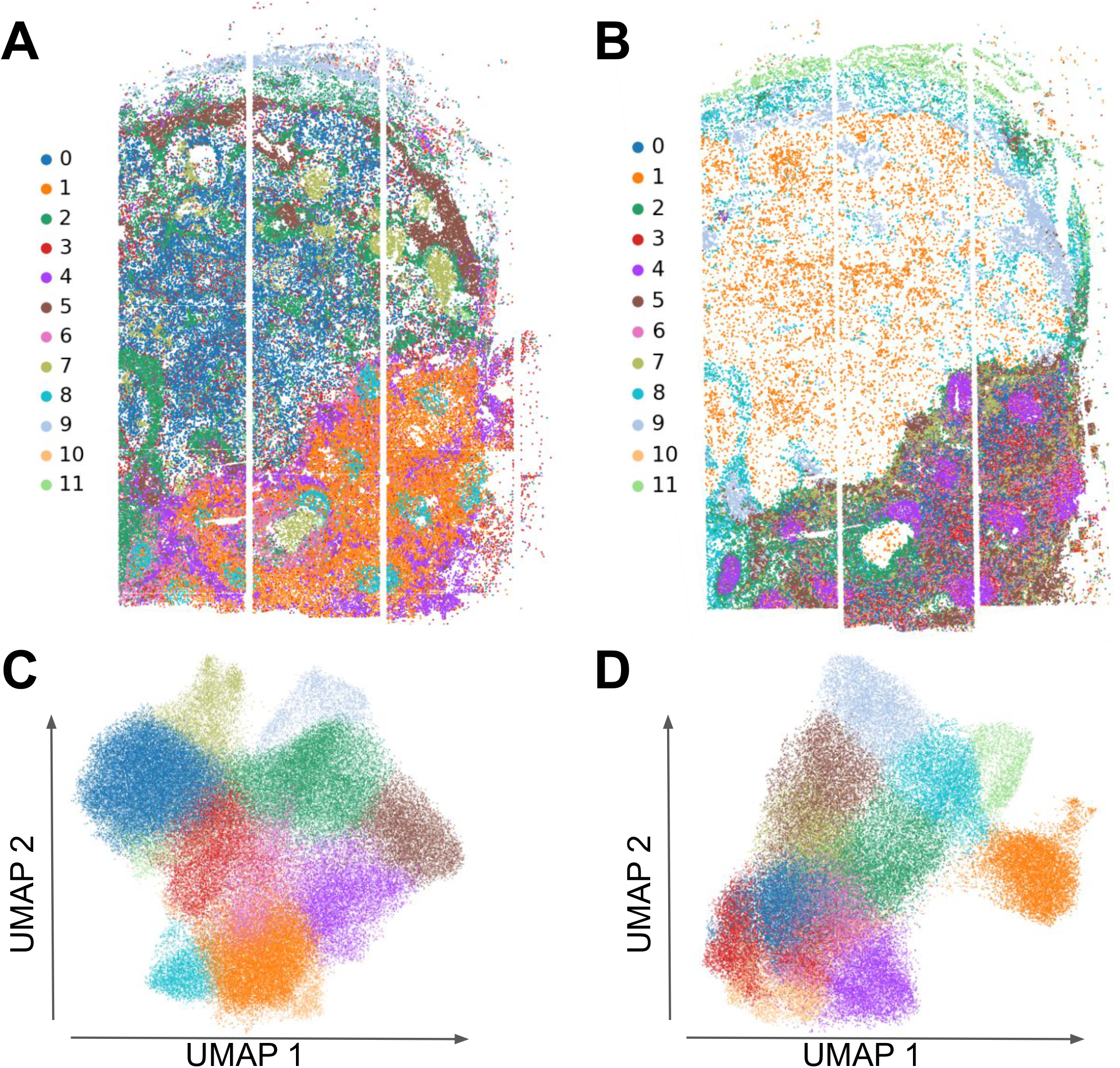
Analysis of sample GSM8990102 (metastatic lymphnode section #4) using FaST (panels A and C) compared to GEO deposited processed data from the same sample. Panel A: Spatial gene expression was reconstructed using FaST. Segmented gene expression in anndata format was used to cluster “pseudo-cells” using scanpy with default parameters. Clustered cells were plotted in spatial coordinates. Panel B: Segmented gene expression in anndata format (GEO) was used to cluster cells using scanpy with default parameters. Clustered cells were plotted in spatial coordinate. Panel C: UMAP plot of pseudo-cells obtained with FaST. Panel D: UMAP plot of segmented cells from GEO processed data (GSM8990102). Cluster colors do not correspond to the same cell types in the two different protocols. Colors were automatically assigned based on the abundance of each cluster in each output (FaST, panels A and C; Spacemake/OpenST panels B and D).

### RNA segmented pseudo-cells recapitulate the tissue composition observed with H&E segmentation

FaST relies on RNA based segmentation which yields “pseudo cells”. To assess the reliability of this method we looked at the clustering of the pseudo-cells as compared to the clustering of the H&E segmented cells, as obtained by the OpenST pipeline. The number of pseudo cells obtained by FaST is roughly comparable to the one obtained by H&E segmentation on both datasets tested. Importantly, the spatial organization of the cell clusters (obtained on scanpy, using default parameters) highlights that, with equal parameters, pseudo-cells yield a slightly larger number of clusters compared to H&E segmented cells. The spatial organization of the clusters observed with FaST mirrors the spatial organization of the clusters observed with OpenST, strongly suggesting that RNA segmented pseudo cells are a very close proxy for H&E segmented cells.

I next asked whether RNA based segmentation is prone to possible artifacts. A major concern in the field of ST cell segmentation is the generation of segmented cells consisting of fragments from different neighboring cells. To assess this issue, we took a set of pairs of genes whose expression is expected to be mutually exclusive as their expression is restricted to different lineages or cell types. We chose CD3E (a marker of T-cells) vs KRT6a (a marker of skin epithelial cells) and ACTA2 (selectively expressed in smooth muscles) vs EPCAM (a marker of epithelial cells). The metastatic lymph node provides a unique tissue in which the cell types expressing these markers are expected to frequently occur in close proximity, and thus provides an excellent model to test for segmentation artifacts.

Our results suggest that cells double positive for both CD3E and KRT6A occur at a lower frequency in FaST output (2.99%) compared to spacemake/OpenST output (8.16%).

On the other hand, cells double positive for ACTA2 and EPCAM occur at a frequency of 1.89% in FaST output, as opposed to a frequency of 1.61% in spacemake/OpenST output (Fig 4). As a ground truth comparison, we are reporting the observed double CD19/CD3E positive cells observed in the “pbmc” scRNA-seq dataset (1.36%) (Figure S4). These data suggest that both H&E and RNA segmentation are very close to the “ground truth” benchmark represented by double CD19/CD3E positive cells in scRNA-seq, and that FaST output does not contain a fraction of inappropriately segmented cells higher that the one obtained with H&E staining based cell segmentation.

**Figure 4:**
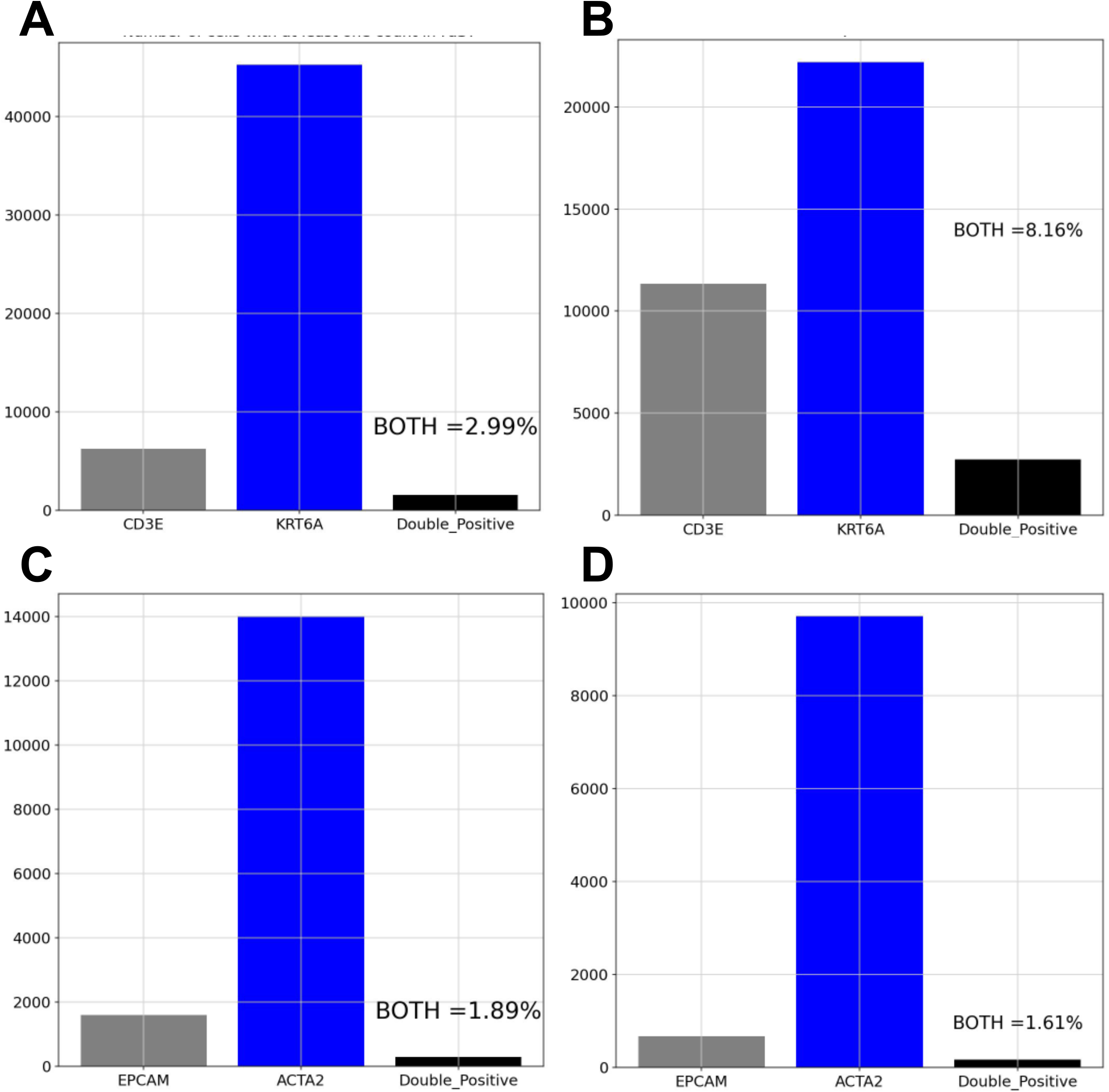
Comparison of the rate cells positives for two marker genes specific to distinct cell lineages in FaST (left panels) and in GEO deposited analysed data (GSM8990102). Panel A: Double positive cells for CD3E (T-cells marker) and KRT6A (epithelial marker) in FaST analysed data. Panel B: Double positive cells for CD3E (T-cells marker) and KRT6A (epithelial marker) in analyzed data from GEO. Panel C: Double positive cells for ACTA2 (Smooth muscle marker) and EPCAM (epithelial marker) in FaST analysed data. Panel D: Double positive cells for ACTA2 (Smooth muscle marker) and EPCAM (epithelial marker) in analysed data from GEO.

## Discussion

The method I am reporting inhere allows fast analysis of Spatially Resolved Transcriptomics datasets corresponding to a tissue section with an area of about 15 mm^2^ in about 2 hours, significantly outperforming existing softwares both in terms of time and memory requirements, thus allowing analysis of such datasets using a small workstation.

Cell segmentation in the context of ST is still an active field of research, with several different approaches proposed thus far. It is worth mentioning inhere that even tissue staining cell segmentation algorithms mainly aim at automating an otherwise very low-throughput task, yet all imaging based cell segmentation algorithms have a tradeoff in terms of accuracy. This is clearly reported in independent systematic comparative assessments of the currently available state of the art methods for staining-driven cell segmentation [20,21]. These studies clearly show that accuracy of these methods is generally good and certainly very useful, but still not identical to the “ground truth”.

The FaST pipeline relies on the spateo-release package to perform a quick and informative cell segmentation based on RNA molecule mappings. The data presented inhere show that, although a perfect overlap between H&E guided and RNA based segmentation is not observed, RNA segmentation faithfully recapitulates the cell types and localizations observed with image-based cell segmentation. The data I report also rule out the possibility that FaST might be more prone to “under-segmentation” artifacts (as assessed by counting cells double positive for incompatible marker genes) compared to existing softwares.

Based on these considerations, I foresee that, by relieving the requirement for tissue staining and image capture and analysis, RNA based segmentation might soon attain sufficient accuracy to replace image based segmentation in most ST data analyses.

Indeed, RNA segmentation might have specific advantages in high resolution ST, as RNA segmented cells share the same coordinates of the barcodes naturally.

FaST pipeline however is modular, thus allowing users to only run mapping and DGE steps and afterwards continue analysis using the preferred (either staining-based or RNA-based) cell segmentation model instead of spateo-release.

Currently, FaST is limited to the ST protocols relying on Illumina flow cells as RNA capture devices. Possible future improvements may include extension to other ST platforms with submicrometer resolution.

FaST was not specifically designed to yield high performance in HPC environments (where the single processors generally have a lower performance compared to the processors of small workstations), however, given that in ST analysis setting the chronological sequence of the different steps of the analysis is relevant, and FaST takes advantage of parallelization whenever possible, my benchmarks confirm that FaST is in fact faster than other existing HPC designed ST analysis softwares also in an HPC computing environment.

## Supplementary Figures

**Figure S1:** Time benchmarking of FaST (either run on a small workstation or in a HPC environment) Spacemake and NovaScope on the different dataset listed in Table 1 and Table 3.

**Figure S2:** Memory footprint of FaST (either run on a small workstation or in a HPC environment), Spacemake and NovaScope on the different dataset listed in Table 1 and Table 3.

**Figure S3:** spatial pattern of expression of 4 marker genes whose expression is confined to specific cell types (KRT6A, epithelial; IGKC, B-cells; FDCSP, Follicular Dendritic cells; CD3E, T-cells), as depicted by the FaST and the spacemake/OpenST pipelines.

**Figure S4:** Assessment of the CD3E/CD19 double positive cells in the “pbmc_3k” dataset.

