## Supplementary Table 1 for "Fast analysis of Spatial Transcriptomics (FaST): an ultra lightweight and fast pipeline for the analysis of high resolution spatial transcriptomics"

**Supplementary Table 1: Benchmarking data for each run of FaST (mapping + SGE)***

| Sample | Reads (M) | Wall time (h:mm:ss) | User time (seconds) | %CPU | RAM (max, Gb) |
|---|---|---|---|---|---|
| Mouse head e13 | 569.6 | 0:43:39 | 23614.97 | 802% | 18.1 |
| Metastatic Lymphnode S4 | 782.7 | 1:12:31 | 53507.34 | 1262% | 18.3 |
| NovaScope N3_B08C_v2 b+bu | 922.1 | 1:27:54 | 43153.52 | 849% | 18.1 |
| NovaScope N3_B08C_v2 b+bu+cu | 1353.9 | 2:12:08 | 67576.88 | 884% | 22.7 |
| NovaScope N3_B08C_v2 b+bu+c+cu | 1781.9 | 2:49:25 | 94788.68 | 965% | 27.8 |

* All tests were carried out on a workstation equipped with an 12th Gen Intel® Core™ i9-12900K and 32Gb of RAM
