## Supplementary figures and images for "Fast analysis of Spatial Transcriptomics (FaST): an ultra lightweight and fast pipeline for the analysis of high resolution spatial transcriptomics"

### Figure S1

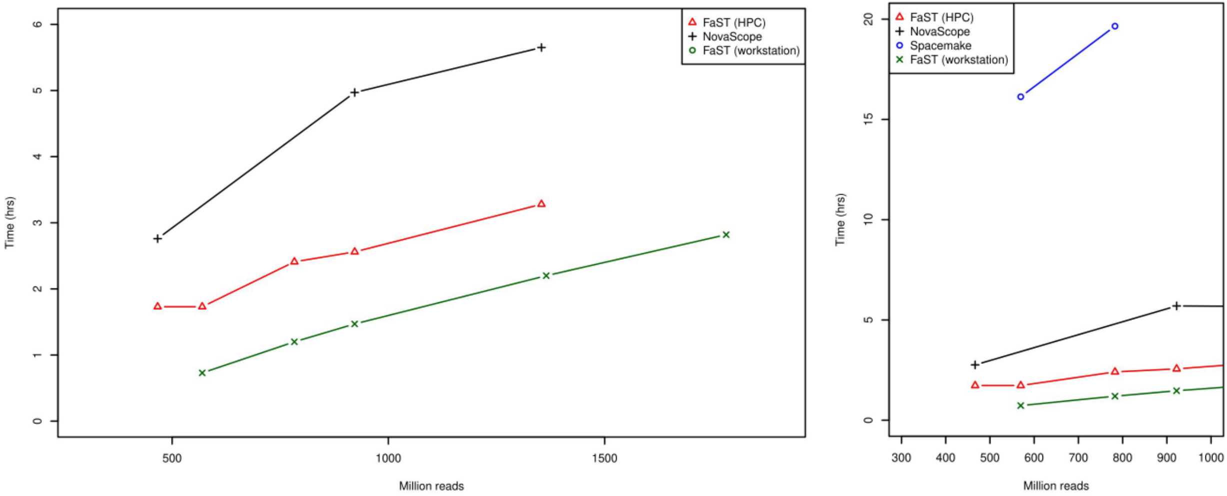

### Figure S2

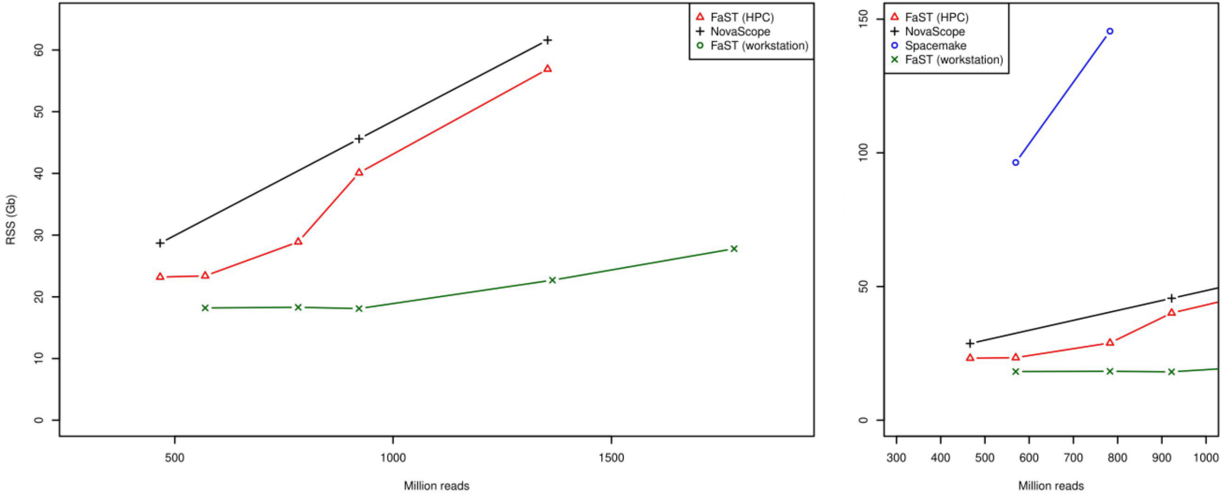

### Figure S3

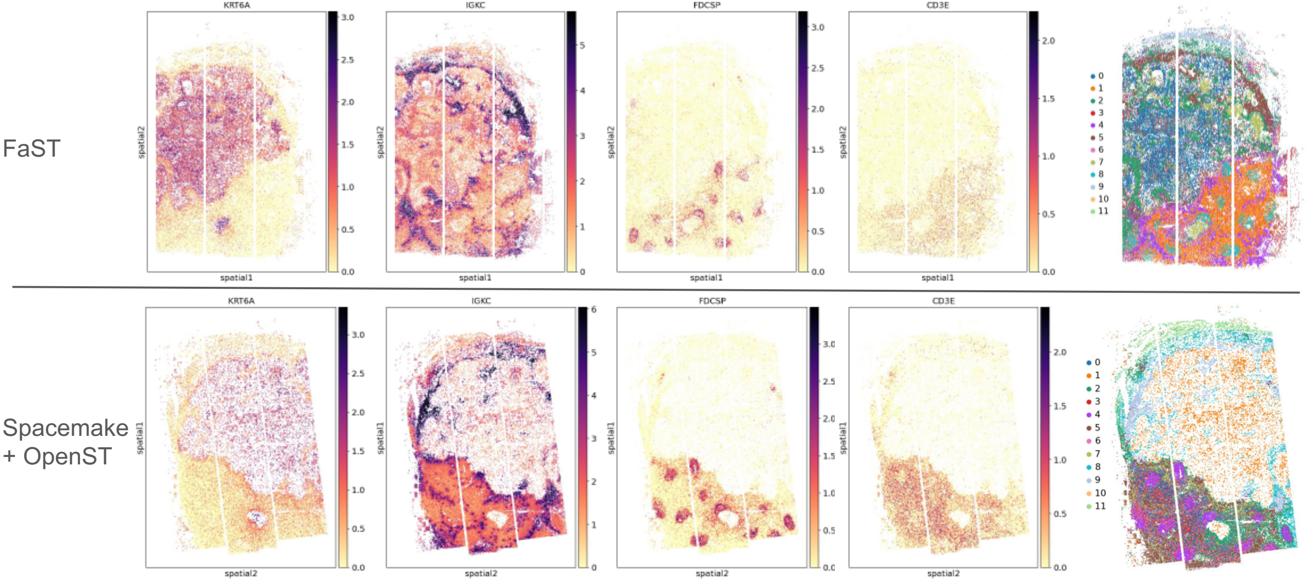

### Figure S4

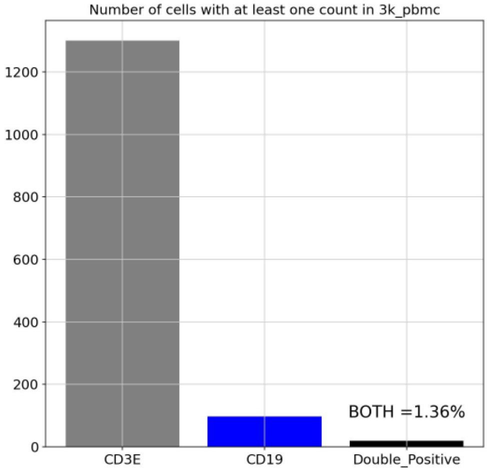
